# Comparing synthetic refocusing to deconvolution for the extraction of neuronal calcium transients from light-fields

**DOI:** 10.1101/2020.09.07.285585

**Authors:** Carmel L. Howe, Peter Quicke, Pingfan Song, Herman Verinaz Jadan, Pier Luigi Dragotti, Amanda J. Foust

## Abstract

**Significance:** Light-field microscopy (LFM) enables fast, light-efficient, volumetric imaging of neuronal activity with calcium indicators. Calcium transients differ in temporal signal-to-noise ratio (tSNR) and spatial confinement when extracted from volumes reconstructed by different algorithms.

**Aim:** We evaluated the capabilities and limitations of two light-field reconstruction algorithms for calcium fluorescence imaging.

**Approach:** We acquired light-field image series from neurons either bulk-labeled or filled intracellularly with the red-emitting calcium dye CaSiR-1 in acute mouse brain slices. We compared the tSNR and spatial confinement of calcium signals extracted from volumes reconstructed with synthetic refocusing and Richardson-Lucy 3D deconvolution with and without total variation regularization.

**Results:** Both synthetic refocusing and Richardson-Lucy deconvolution resolved calcium signals from single cells and neuronal dendrites in three dimensions. Increasing deconvolution iteration number improved spatial confinement but reduced tSNR compared to synthetic refocusing. Volumetric light-field imaging did not decrease calcium signal tSNR compared to interleaved, widefield image series acquired in matched planes.

**Conclusions:** LFM enables high-volume rate, volumetric imaging of calcium transients in single cells (bulk-labeled), somata and dendrites (intracellular loaded). The trade-offs identified for tSNR, spatial confinement, and computational cost indicate which of synthetic refocusing or deconvolution can better realize the scientific requirements of future LFM calcium imaging applications.

## 1 Introduction

Understanding how neuronal networks learn, process, and store information requires imaging techniques capable of monitoring the activity of hundreds to thousands of neurons simultaneously in three-dimensional (3D) tissues. Capturing rapid neuronal calcium dynamics requires high temporal resolution at cellular or subcellular spatial resolution.^1^ The development of synthetic and genetically-encoded fluorescent indicators of intracellular calcium concentration^2, 3^ and membrane voltage^4, 5^ enables functional imaging on these scales.

The optical sectioning capability of confocal and multi-photon scanning microscopes adapts them well to 3D imaging of scattering brain tissues. However, scanning limits the fluorescence bandwidth and hence the acquisition speed and temporal signal-to-noise ratio (tSNR). tSNR describes the ability to discriminate transient changes in fluorescence from baseline noise. For shot noise-limited systems, tSNR is proportional to the square-root of the collected fluorescence photon flux. That is why applications requiring high acquisition rates and/or SNR typically rely on wide-field, single-photon imaging to maximize photon flux by exciting fluorescence simultaneously in all illuminated structures. Widefield excites fluorescence efficiently throughout a volume, however, only one axial plane is imaged. In this configuration, fluorescence excited above and below the imaging plane is not only unnecessary, but contributes spurious fluorescence to the in-focus image, degrading contrast and confusing the functional signals.^6^

Light-field microscopy (LFM) exploits out-of-focus fluorescence simultaneously excited throughout the volume. LFM combined with widefield, single-photon fluorescence excitation enables volumetric collection, maximizing the photon budget. LFM is a 3D imaging technique, which encodes both lateral position and angular information, unlike conventional imaging that focuses on objects in a single plane.^7^ A microlens array (MLA) at the microscope’s native image plane enables image reconstruction at different planes and perspectives from a single light-field image. This increases light efficiency and speed at the cost of spatial resolution as the cameras pixels now divide over four-dimensions (x,y,*θ*_*x*_,*θ*_*y*_) rather than two (x,y). The four-dimensional light-field can be used to reconstruct a volume around the native focal plane, slice by slice. Two methods for reconstructing volumes from LFM images are commonly used: synthetic refocusing^7^ and 3D deconvolution.^8^ Synthetic refocusing extracts single planes from a light-field that correspond to widefield images. Multiple planes can be reconstructed orthogonal to the optical axis to generate a *z*-stack. Synthetic refocusing is computationally fast as each pixel in the output volume is simply the weighted sum of a subset of pixels in the light-field. However, similar to widefield imaging, this technique lacks optical sectioning such that out-of-focus sources reduce the contrast of in-focus sources. In contrast, 3D deconvolution reconstructs a volume by deconvolving its light-field measurements with a 3D light-field Point Spread Function (PSF) based on a wave optics model^9^ of the LFM. This can be achieved by using iterative deconvolution methods, such as the Richardson-Lucy^10, 11^ or Image Space Reconstruction Algorithms.^12^ 3D deconvolution can achieve a higher spatial resolution than synthetic refocusing because the individual projections through the volume sample the object more finely than the microlens array, thus improving the discrimibility of signals in 3-dimensions. However, 3D deconvolution approaches are computationally intensive and amplify noise.^13^

LFM’s capacity to capture volumetric data from 2D frames has recently motivated its application to imaging neuronal activity in non-scattering specimens such as C. Elegans and Zebrafish,^14–19^ and in mammalian brain *in vivo*.^20–22^ Seeded iterative demixing^20, 22^ and compressive LFM^15^ increase the speed of neuronal localization and single-cell time series analysis by identifying and localizing somatic signals. Notably, these techniques improved performance in scattering brain tissues compared to volume reconstruction methods that only account for ballistic photons. However, volume reconstruction is still necessary to image the generation and propagation of voltage and calcium transients spatially extended structures such as axons and dendrites.

Here we show that LFM can resolve calcium transients simultaneously in axially separated somata and dendrites of neurons loaded with a red-emitting calcium dye, CaSiR-1.^23^ We examined trade-offs between the tSNR and the spatial signal confinement of calcium signals localized in volumes reconstruction from light fields by synthetic refocusing and 3D deconvolution. A comparison of calcium signals extracted from interleaved light-field and widefield imaging trials showed no penalty to tSNR for light-field trials, which additionally enabled localization of calcium signals in 3D. These results demonstrate the power of LFM for simultaneously tracking calcium transients in axially separated neurons and neuronal subcompartments. By distilling the trade-offs between spatial signal confinement and tSNR, these results underline the importance of selecting a volume reconstruction method adapted to the scientific goals of future experiments.

## 2 Materials and Methods

Parts of the following methods and preliminary SNR quantification results were published in Howe *et al*. (2020).^24^

### 2.1 Optical System

We designed our LFM following Levoy *et al*. (2006).^7^ Imaging was performed with a custom-built epifluorescence microscope with a MLA (125 µm pitch, f/10, RPC Photonics) placed at the imaging plane of a 25×, Numerical Aperture (NA)=1.0 water immersion objective lens (XLPLN25XSVMP, Olympus) and 180 mm tube lens (TTL180-A, Thorlabs), illustrated in Figure 1A. The MLA was imaged onto a scientific complementary metal-oxide-semiconductor (sCMOS) camera (ORCA Flash 4 V2 with Camera Link, 2048×2048 pixels, 6.5 µm pixel size, Hamamatsu) with a 1:1 relay macro lens (Nikon 60 mm f2.8 D AF Micro Nikkor Lens).

**Fig 1.**
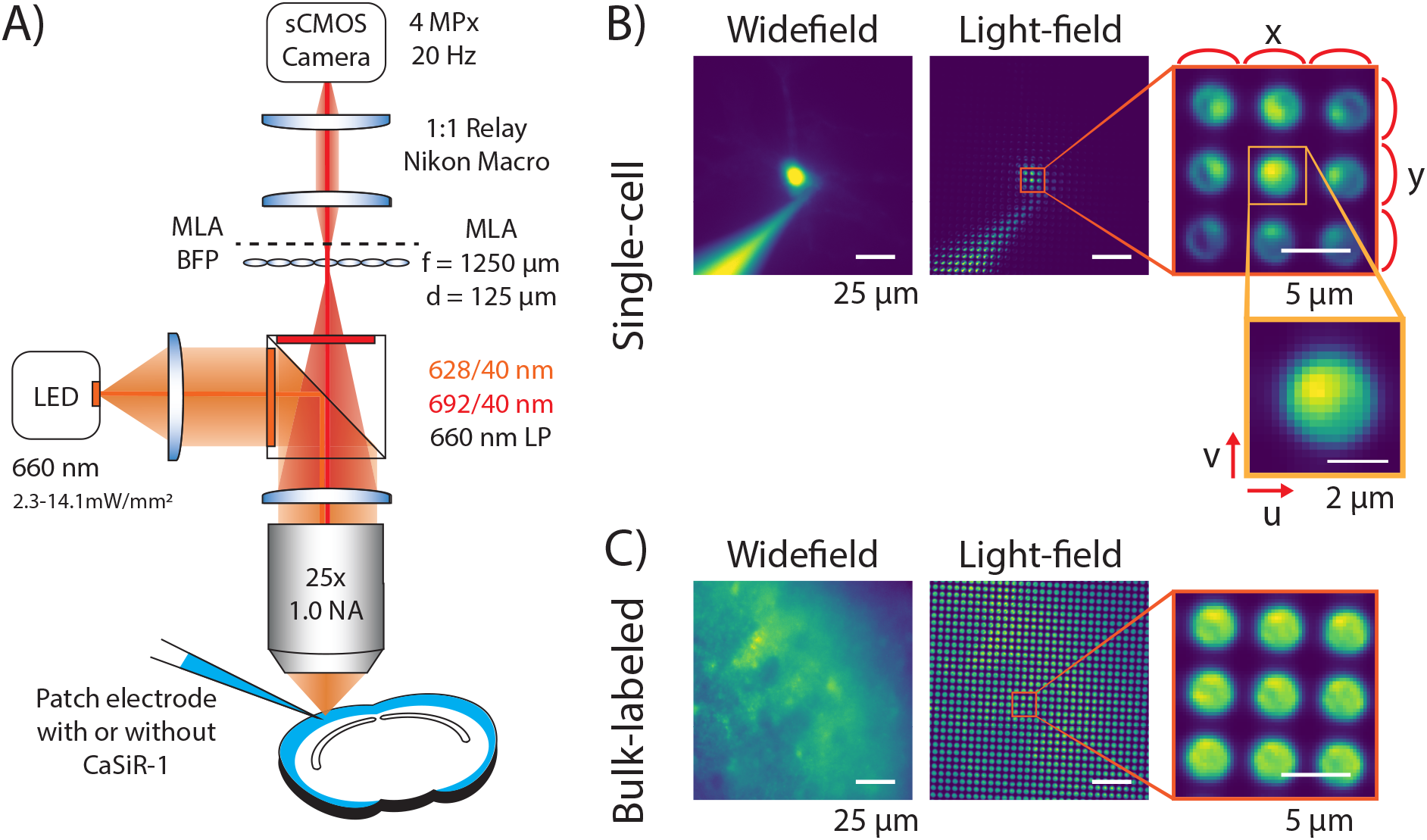
A) Optical system schematic. A microlens array is placed at the native imaging plane of a widefield microscope and the back focal plane is imaged onto a sCMOS camera enabling 3D reconstructions from a 2D frame. B) Example widefield and light-field images from both a single neuron intracellularly loaded with the synthetic calcium dye, CaSiR-1 via a micropipette. Close-up views of the raw light-field images show the circular subimages encoding the 4D spatial and angular information. The light-field is parameterized by a 4D function, ℒ (*u, v, x, y*), where each lenslet is ℒ (*u, v*, ·, ·) and the same pixel in each lenslet subimage is ℒ (·, ·, *x, y*). C) Example images from bulk-labeled slices where CaSiR-1 AM was bath applied to many neurons.

The LFM image consists of circular subimages (Figure 1B) which are parameterized by the 4D function ℒ (*u, v, x, y*), where each lenslet is ℒ (*u, v*, ·, ·) and the same pixel in each lenslet subimage is ℒ (·, ·, *x, y*). Each circular subimage represents the angular content of the light at a specific spatial location.

The ‘native LFM spatial resolution’ is given by the microlens pitch divided by the objective magnification. Therefore, an MLA was chosen such that the lateral resolution of our LFM was 5 µm, roughly half the diameter of a cortical neuron (10 µm). The axial resolution of a LFM is defined by the number of resolvable diffraction-limited spots behind each microlens.^7^ Using the Sparrow criterion and assuming a peak emission wavelength of 664 nm (*λ*) for CaSiR-1,^23^ the spot size in the camera plane is 7.64 µm. So, with a 125 µm pitch MLA, we are able to resolve *N*_*u*_ = 13 distinct spots under each microlens. The depth of field when synthetically refocusing is given by eq. (1), resulting in a depth of field of 6.52 µm^7^ compared to 0.8 µm in a conventional widefield microscope with the same imaging parameters.

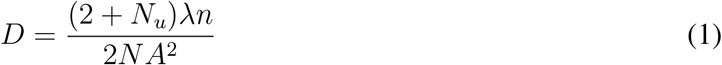

where *n* is the refractive index.

### 2.2 Brain slice preparation

This study was carried out in accordance with the recommendations of the UK Animals (Scientific Procedures) Act 1986 under Home Office Project and Personal Licenses (project license 70/9095). 400 µm slices were prepared from 33 to 196 day old mice using the ‘protective recovery’ method.^25^ Slices were cut in Na-aCSF containing (in mM): 125 NaCl, 25 NaHCO_3_, 20 glucose, 2.5 KCl, 1.25 NaH_2_PO_4_, 2 MgCl_2_, 2 CaCl_2_. After cutting, the slices were transferred for a period of 12 minutes to a solution containing (in mM) 110 N-Methyl-D glucamine, 2.5 KCl, 1.2 NaH_2_PO_4_, 25 NaHCO_3_, 25 Glucose, 10 MgCl_2_, 0.5 CaCl_2_, adjusted to 300 – 310 mOsm/kg, pH 7.3 – 7.4 with HCl at 36^°^C, before being transferred back to the first solution for at least an hour before imaging trials. All solutions were oxygenated with 95% O_2_/5% CO_2_.

After resting the slices were either bulk-labeled with CaSiR-1 AM-ester dye or used for single-cell labeling with CaSiR-1 potassium salt.

For bulk-labeled slices 50 µg, CaSiR-1 AM (GC402, Goryo Chemicals)^23^ was dissolved in 10 µl of dimethyl sulfoxide (DMSO) with 10% w/v Pluronic F-127 (Invitrogen) and 0.5% v/v Kolliphor EL (Sigma-Aldrich).^26^ The slices were then incubated for 40 minutes at 37^°^C in 2 ml of Na-aCSF with the CaSiR-1 AM/DMSO mixture pipetted onto the surface of each slice, oxygenated by blowing 95% O_2_/5% CO_2_ onto the surface. After loading, the slices rested in room temperature Na-aCSF for at least 20 minutes before use.

### 2.3 Imaging

For single-cell labeling, cortical cells were patched using 6 – 8 MOhm patch pipettes containing intracellular solution consisting of (in mM): 130 K-Gluconate, 7 KCl, 4 ATP-Mg, 0.3 GTP-Na, 10 Phosphocreatine-Na, 10 HEPES, 0.1 CaSiR-1 potassium salt (GC401, Goryo Chemicals).^23^ After sealing and breaking in, the calcium dye was allowed to diffuse into the cell (Figure 1B). For bulk-labeled slices (Figure 1C), cortical cells were patched containing the same intracelluar solution without the addition of the CaSiR-1 potassium salt.

Cells were patched under oblique light-emitting diode (LED) infrared illumination (peak 850 nm). The signals were recorded with a Multiclamp 700B amplifier (Axon Instruments) and digitized with a Power 1401 (Cambridge Electronic Design).

Imaging trials were taken at 20 frames/s at room temperature. Stimulation consisted of five current pulses for 10 ms at 0.5 Hz where the current was adjusted to stimulate a single action potential. For single cells, this stimulus was applied to the labeled cell with the dye-loading pipette. For bulk-labeled slices, the stimulus was applied to a cell in the field of view causing broader activation of multiple neurons in the local network. Widefield and light-field trials were interleaved by removing and replacing the MLA from a precision magnetic mount (CP44F, Thorlabs). The removal and addition of the MLA shifted the focal sample plane. We calculated this focal plane shift using the thin lens equation to be ±2 µm.

Fluorescence was excited with a 660 nm LED (M660L2, Thorlabs) powered by a constant current source (Keithley Sourcemeter 1401) to illuminate the sample between 2.3-14.1 mW/mm^2^. The 660 nm LED was collimated with an f = 16 mm aspheric lens (ACL25416U0-A, Thorlabs) and filtered with a 628/40 nm excitation filter (FF02-628/40, Semrock). Collected fluorescence was filtered with a 660 nm long-pass dichroic (FF660-Di02, Semrock) along with a 692/40 nm emission filter (FF01-692/40, Semrock). Imaging data were acquired with Micromanager.^27^

Single-cell labeled somata laid between 46 and 49 µm below the slice surface, with a median depth of 47 [IQR, 46.2, 48.6] µm. Whereas bulk-labeled somata were between 29 and 36 µm below the slice surface, with a median depth of 34 [30, 34.8] µm.

### 2.4 Light-field volume reconstruction

We reconstructed light-field source volumes from the raw light-fields (Figure1B&C) using synthetic refocusing^7^ and Richardson-Lucy (RL) 3D deconvolution.^8, 10, 11^ Images synthetically refocused at a plane *f′* = *αf*_0_, where *f*_0_ is the native focal plane, were calculated from a light-field image parameterized by ℒ (*x, y, u, v*) using the formula derived in^28^ as

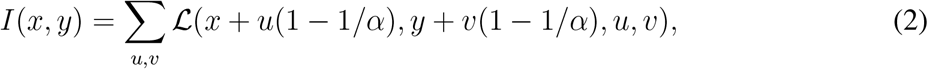

where *I*(*x, y*) represents the refocused image. This process can be interpreted as a summation over different shifted angular ‘views’ of the sample represented by ℒ (*x, y*, ·, ·) such that the rays forming the views intersect at the desired refocus plane. We synthetically refocused ‘stacks’ of images or image time series, *I*(*x, y, z, t*) at 1 µm *z*-intervals using linear interpolation of the collected light-field images or videos.

Stacks from the same light-field images were also calculated using RL deconvolution. The 3D light-field PSF was calculated using the method described in,^9^ by considering how a LFM collects fluorescence from a dipole oscillating with a wavelength of 550 nm. The total PSF was calculated as an incoherent sum of dipoles oriented along *x, y*, and *z*. PSF values were calculated on a 5 × 5 grid relative to the microlens. A low resolution PSF was calculated by averaging over the PSF values weighted by a 2D Hamming window of a width equal to the MLA pitch and coaxial with the lens. The estimated volume, *x*, is recovered from the measured light-field image, *y*, and the PSF, *H* using the following iterative update scheme in matrix-vector notation:

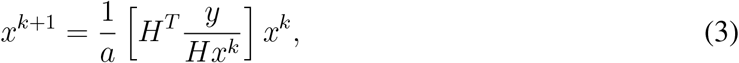

where the fraction *y/Hx*^*k*^ is computed element-wise and *a* = ∑_*i*_ *H*(*i*, :). Stacks were reconstructed using this method as with synthetic refocusing for varying numbers of iterations of eq. 3.

Additionally, to enhance edges and reduce noise, we slightly modified the objective function of RL to include a total variation (TV) term, as in.^29^ To incorporate this regularization prior, we modified the standard RL as follows

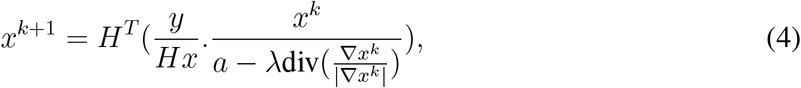

where div is the divergence operator, ∇ is the gradient operator, and *λ* is a regularization factor set to 0.01, determined by visual inspection of the volumes.

### 2.5 Time Series Analysis

#### 2.5.1 SNR

Signals were extracted from widefield or light-field time series reconstructed with synthetic refocusing or RL 3D deconvolution.

We calculated Δ*F/F* using eq. (5) where *F* was the raw fluorescent signal, *F*_0_ was the baseline fluorescence taken as an average prior to the action potential, and *F*_*d*_ was the camera’s dark signal (all in counts).

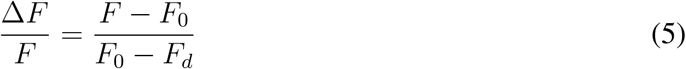

An ‘activation map’ was produced from the variance over time to indicate the pixels containing the greatest temporal signal from the Δ*F/F* map. Regions of interest (ROI) were defined by extracting the top 2 percentile of signal containing pixels (somatic and dendritic).

The SNR was calculated by dividing the peak signal (%) by the baseline noise (%), given by the square-root of the variance of the baseline fluorescence taken as an average prior to the action potential (20 samples, 1 second).

### 2.6 Statistics

All statistics are reported as median [inter-quartile range (IQR)]. Wilcoxon matched-pairs signed-rank test was performed between synthetically refocused and 3D deconvolved light-field time series. These reconstructions were generated from the same image series, removing independent variables such as bleaching and changes in dye loading in the case of single-cell labeling. Statistical analysis was performed using Python SciPy.^30^

### 2.7 Signal Confinement

#### 2.7.1 Spatial Profiles

To compare the signal confinement spatial profiles were generated. To produce the widefield axial profile a *z*-stack was collected manually. At the end of an imaging trial the micropipette was removed and a *z*-stack was collected by moving the plane of focus through the sample between -40 to 40 µm in steps of 1 µm using a stepper motor. Light-field axial profiles (*xz, yz*) were generated by synthetically refocusing and deconvolving at different depths of focus from the light-field taken with the cell in the native imaging plane. Lateral profiles (*xy*) were then generated by taking a line plot through the cell on widefield or reconstructed light-field images at the plane of best focus. The spatial signal confinement is reported from the Full-Width at Half-Maximum (FWHM). Friedman’s Two-Way Analysis of Variance by Ranks was performed between the FWHMs from widefield and light-field volumes reconstructed with synthetic refocusing and 3-iteration RL 3D deconvolution.

The spatial profiles from single-cells were generated from either a single static image in the case of light-field frames or a stack of widefield frames. However, in bulk-labeled slices the background signal was very large and the spatial profiles were generated from the activation map described in Section 3. Maximum intensity projections were taken through *xy, xz*, and *yz*.

#### 2.7.2 Temporal Spatial Profiles

Temporal spatial profiles were produced from single cells to determine the axial spread of the calcium fluorescence response. Light-field axial profiles were generated as in Section 2.7.1. Time courses were extracted for each depth from either a somatic or nearby dendritic ROI. Δ*F/F* was calculated using eq. 5 in Section 2.5.1. A line plot across the axial range was generated from the sum over time.

## 3 Results

### 3.1 Synthetic refocusing enables fast, high SNR light-field reconstruction

We compared the performance of light-field reconstruction techniques on the tSNR of CaSIR-1 signals extracted from both single-cell (intracellularly loaded) and bulk-labeled slices. We reconstructed volumetric light-field time series from 4 single cells (Figure 2A & Supplementary videos S2A & B) and 4 bulk-labeled slices (Figure 2B) with synthetic refocusing and Richardson-Lucy 3D deconvolution. For single-cell trials, calcium transients were stimulated by applying suprathresh-old current pulses (red lines) to the soma in whole-cell current clamp (Figure 2A2). Calcium transients from bulk-labeled slices were captured after a single cell was stimulated within the field of view (Figure 2A2). We interleaved widefield and light-field acquisitions to facilitate comparison of functional signals extracted from matched ROIs. Time courses were extracted from a ROI taken from the top 2 percentile of pixels at the native focal plane. The SNR, peak signal, and baseline noise were compared between the two light-field reconstruction algorithms and widefield image series.

**Fig 2.**
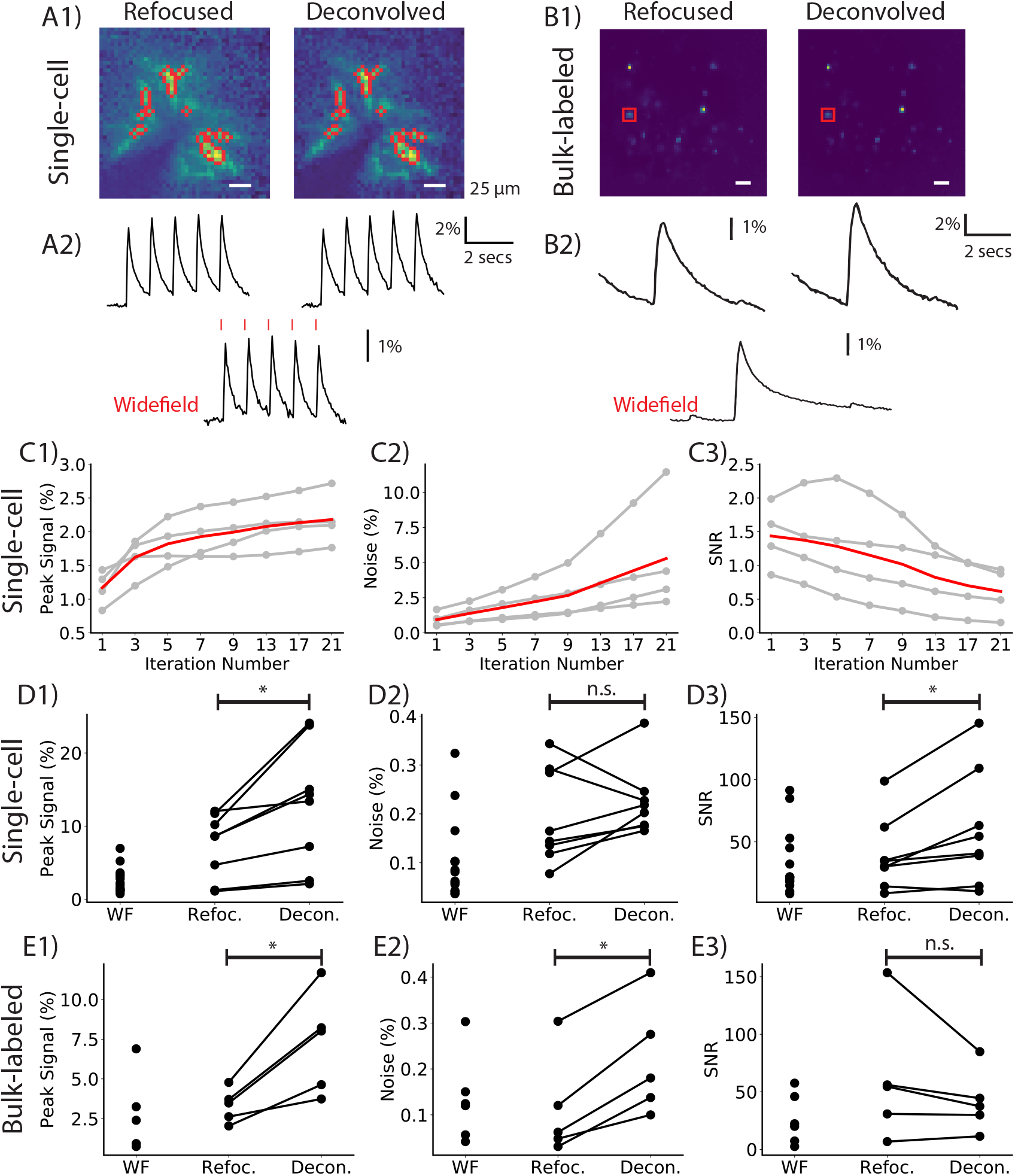
A comparison of calcium transient tSNR for widefield (WF) and light fields reconstructed with temporal refocusing or RL 3D deconvolution. A1 and B1 show the calcium activation maps of planes reconstructed from light fields containing a single labeled cell (A1) and multiple cells in bulk-labeled slices (B1) using synthetic refocusing and RL 3D deconvolution (3-iteration) algorithms. Calcium transient time series (A2,B2) were extracted from the mean pixel intensities of the ROIs (outlined in red). As deconvolution iteration number increases, so does the peak signal (C1) and noise (C2) respective to time series reconstructed with synthetic refocusing for matching ROIs, ultimately reducing the SNR (C3). The gray traces are from separate single-cell experiments and the red line is the average (n=4 cells). C1-C3 are normalized by the signal, noise, and SNR of signals extracted from the same ROIs in the synthetically refocused planes. D and E compare peak signal (%), noise (%), and SNR between time series extracted from WF images series, refocused and deconvolved (3-iteration RL) light fields.

Iterative 3D deconvolution algorithms including Richardson-Lucy are known to amplify noise^29^ which increases with iteration number. Therefore, we quantified the effect of iteration number on the peak signal, noise, and SNR from single-cell trials. light-field time series were deconvolved with between 1 and 21 iterations. The deconvolved time series were normalized to synthetically refocused time series generated from the same raw light-fields. On average, the peak signal (%) increases with iteration number with respect to synthetically refocused light-field time series (Figure 2C1). Between 1 and 7 iterations, the deconvolved peak signal increases after which it plateaus with a peak signal around 2× greater than that achieved by synthetic refocusing. In all trials, as iteration number increases, the noise (%) increases compared to synthetically refocused light-field time series (Figure 2C2). The deconvolved time series noise was on average the same as synthetically refocused light-field time series after 1 iteration increasing to 5× greater with 21 iterations. Therefore, on average, as iteration number increases the SNR reduces (Figure 2C3). The SNR from deconvolved light-field time series after 1 iteration is on average 1.5×larger than that of synthetically refocused light-field time series. The SNR from deconvolved and synthetically refocused trials is the same around 9 iterations. Deconvolution tSNR decreases to half that of synthetically refocused after 21 iterations.

Next, we compared the performance of light-field reconstruction techniques on the SNR from all trials for both single-cell and bulk-labeled slices. Three-iteration RL deconvolution was chosen to give the best lateral signal confinement at the highest possible SNR, as detailed in the next section.

The peak signal from single-cell trials (8 trials, 4 cells, 3 mice) was significantly larger when extracted from light-field time series reconstructed with three-iteration Richardson-Lucy 3D deconvolution (13.9 [2.4, 23.9]%) compared to synthetic refocusing (8.6 [1.2, 11.8]%; Wilcoxon matched pairs signed rank, n = 8, w = 36.0, p = 0.01) and single-plane widefield time series (3.2 [1.7, 5.6]%; Figure 2D1). The baseline noise did not differ between light-field time series reconstructed with three-iteration 3D deconvolution (0.21 [0.17, 0.29]%), those reconstructed with synthetic refocusing (0.15 [0.11, 0.31]%; Wilcoxon matched pairs signed rank, n = 8, w = 28.0, p = 0.02), and those from widefield time series (0.10 [0.04, 0.26]%; Figure 2D2). The SNR of times series from three-iteration RL-deconvolved frames (47.6 [13.3, 120.0]) was significantly greater than that of synthetically refocused frames (32.5 [12.6, 73.0]; Wilcoxon rank sum, n = 8, w = 25.0, p = 0.03) and single-plane widefield time series (21.9 [16.8, 86.2]; Figure 2D3).

In bulk-labeled slices (5 trials, 4 cells, 2 mice), the peak signal was significantly greater for light-field time series reconstructed with three-iteration RL 3D deconvolution (8.0 [4.1, 10.3]%) compared to synthetically refocused (3.5 [2.3, 4.4]%; Wilcoxon rank sum, n = 5, w = 15, p = 0.04) and widefield time series (1.7 [0.8, 5.1]%; Figure 2E1). The baseline noise was significantly larger in three-iteration deconvolved bulk-labeled slices (0.18 [0.12, 0.35]%) compared to synthetic refocusing (0.06 [0.04, 0.23]%; Wilcoxon rank sum, n = 5, w = 15, p = 0.04), and widefield time series (0.12 [0.05, 0.22]%; Figure 2E2). The SNR from light-field time series reconstructed with synthetic refocusing (54.5 [16.3, 114.5]) did not differ from deconvolution-reconstructed trials (37.4 [18.7, 68.7]; Wilcoxon rank sum, n = 5, z = 2.0, p = 0.14) or widefield trials (21.1 [4.9, 51.7]), Figure 2E3).

To enhance edges and reduce noise in bulk-labeled volumes, we modified the objective function of RL to include a TV regularization term (Figure S1A). Inclusion of the TV term in the RL deconvolution reduced the total variation of the deconvolved stacks from 0.16 to 0.123 after 10 iterations. However, the mean squared error between TV and non-TV reconstructed volumes was very small, resulting in identical peak signal, noise, and SNR in the extracted calcium time series (Figure S1B). Increasing iteration number up to 30 reduced peak signal, and thus SNR, for the TV-regularized volume (Figure S1C).

### 3.2 Deconvolution reconstruction algorithms provide enhanced spatial signal confinement

We compared the lateral and axial signal confinement of single cells intracellularly labeled with calcium dye between widefield z-stacks and 3D light-fields reconstructed with synthetic refocusing (Figure 3A2) and RL 3D deconvolution (Figure 3A3 & Supplementary video S2C). To assess the impact of deconvolution iteration number on spatial confinement, we measured the FWHM of lateral and axial profiles, normalized to the FWHM the same profiles in synthetically refocused volumes. Both the lateral (Figure 3B1) and axial (Figure 3B2) signal confinement increase with increasing deconvolution iteration number. The red line shows the average for the three cells. The lateral signal confinement (Figure 3B1) for one iteration is 1.6× better than synthetically refocused light-field images and plateaus around 7 iterations with a 2× improvement. The axial signal confinement (Figure 3B2) for one deconvolution iteration is 1.4× better than synthetic refocusing increasing to 2.5× after 21 deconvolution iterations. Three-iteration Richardson-Lucy deconvolution was chosen for further analysis as it maximized lateral confinement while maintaining a high tSNR.

**Fig 3.**
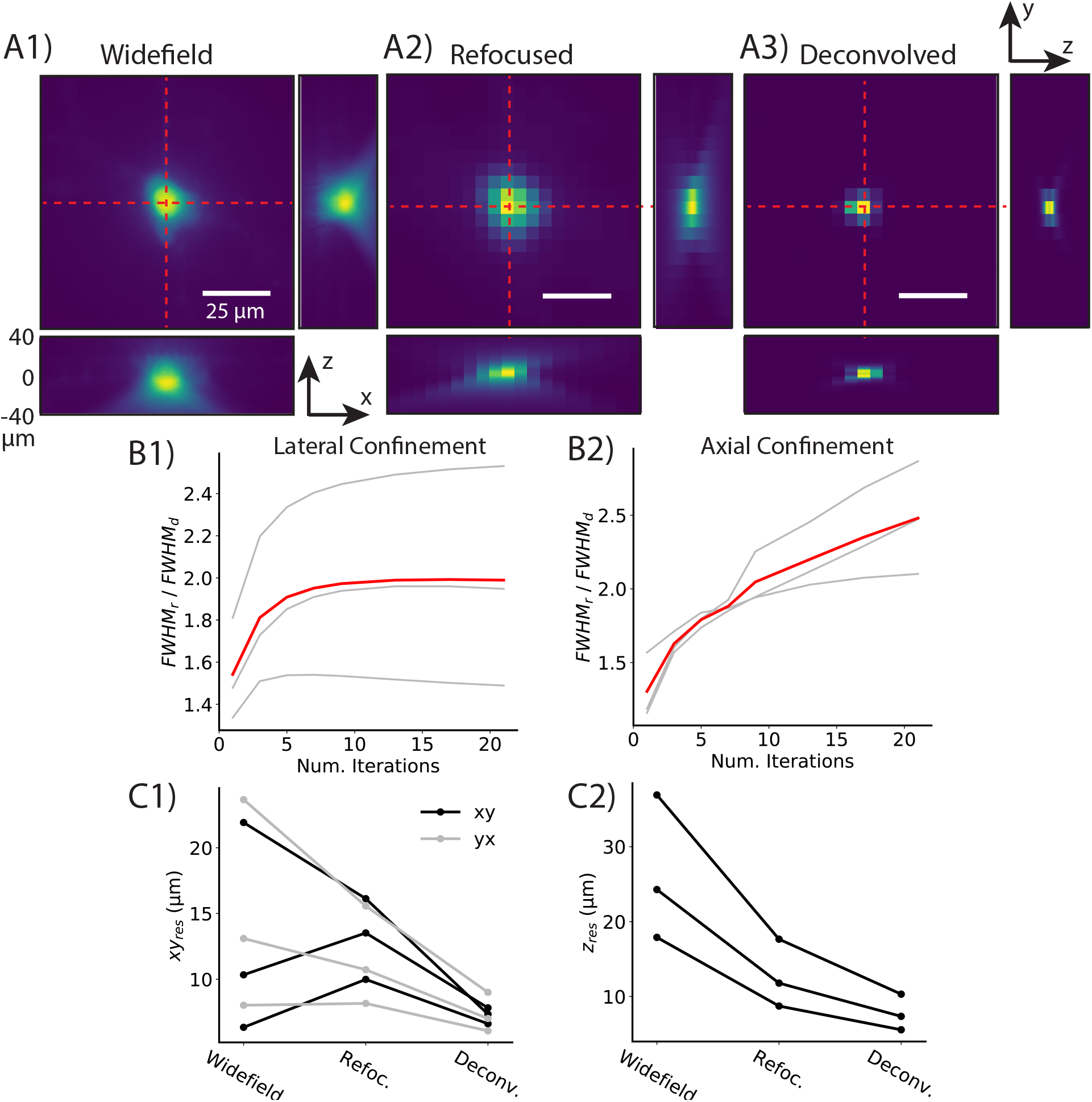
Deconvolution enhances spatial signal confinement compared to widefield stacks and light-field volumes reconstructed with synthetic refocusing. Lateral and axial profiles from a single-cell filled with CaSiR-1 dye are shown. The lateral profiles are plotted at the native focal plane from widefield stacks (A1), and light-field volumes reconstructed with synthetic refocusing (A2) and 3-iteration Richardson-Lucy deconvolution (A3). The axial profiles have been extracted from the lateral position intersected by the red dashed lines at depths ranging from -40 to +40 µm. Increasing deconvolution iteration number increases both the lateral (B1) and axial (B2) signal confinement compared to synthetically refocused volumes. The deconvolved FWHMs are normalized to that of synthetic refocusing. The gray lines are from three different cells, and the red line is the average. Deconvolved light-fields (3-iteration RL) features better lateral (C1) and axial (C2) spatial confinement than widefield z-stacks and synthetically refocused light-field volumes.

The 2D spatial profiles (Figure 3A1-3) clearly show that the light-field images reconstructed with 3D deconvolution have better spatial signal confinement, both laterally and axially compared to both those reconstructed with synthetic refocusing and widefield stacks. The spatial profile for refocused volumes looks similar to widefield, which is expected due to the nature of the reconstruction. A line plot was taken through the lateral and axial profiles, and the FWHM was calculated for each of the imaging configurations from 3 cells (Figure 3C1&2). The results are summarized in Table 1.

**Table 1.** Summary of FWHM from single-cell labeled spatial profiles. Reported as median [IQR], n=3. *3-iteration Richardson-Lucy.

|  | Widefield | Refocused | Deconvolved* |  |
| --- | --- | --- | --- | --- |
| $xy$ | 10.3 [7.1, 19.6] | 13.5 [10.7, 15.6] | 7.3 [6.8, 7.7] | $\mu\text{m}$ |
| $yx$ | 13.1 [9.0, 21.6] | 10.7 [8.7, 14.6] | 7.0 [6.3, 8.6] | $\mu\text{m}$ |
| $xz$ | 17.9 [19.2, 34.4] | 11.8 [9.3, 16.5] | 7.4 [5.9, 9.7] | $\mu\text{m}$ |

The lateral signal confinement (*xy* & *yx*; Figure 3C1) from light-field images reconstructed with 3D deconvolution (3-iteration RL) was not significantly better than that of synthetically refocused or widefield stacks (Friedman’s Two-Way Analysis of Variance by Ranks; *xy*: n=3, w=2.67, p = 0.26 *yx*: n=3, w=4.67, p = 0.10). However, 3D deconvolution significantly improved axial signal confinement (*xz*; Figure 3C2) compared to that of synthetically refocused or widefield stacks (Friedman’s Two-Way Analysis of Variance by Ranks; n=3, w=6, p ¡ 0.05).

For the bulk-labeled slices, the low contrast of the raw images precluded segmentation of individual cells. The cellular spatial profiles were therefore generated from an activation map (the variance over time). Maximum intensity projections through *xz* and *yz* are shown (Figure 4). The signal confinement for both synthetically refocused and 3D deconvolved light-field volumes enabled resolution of a number of active neurons across different focal planes spanning about 9 µm, which is unachievable with any widefield imaging system. The center of mass of each neuron ranges from depths of -5 to +4 µm. The image contrast is higher for 3D deconvolved than for refocused volumes.

**Fig 4.**
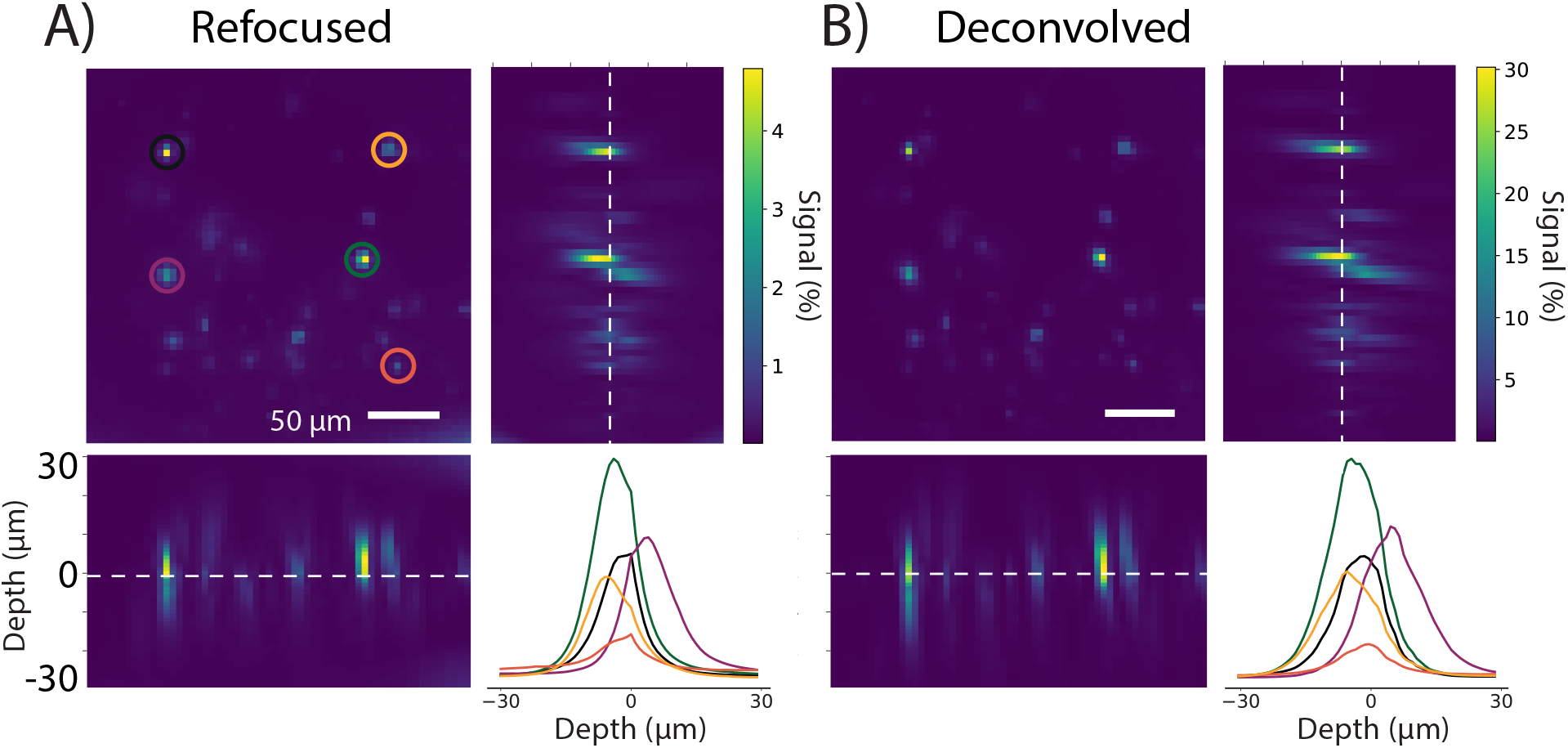
Reconstructed light-field volumes can distinguish cells from different axial planes in bulk-labeled slices. Planes from bulk-labeled slices were reconstructed from light-field volumes with synthetic refocusing (A) and 3D deconvolution (B, 3-iteration Richardson-Lucy) between -30 and +30 µm in steps of 1 µm. An activation map was generated from the variance over time to identify active neurons. A maximum intensity projection through *z* was generated. A *xz* and *yz* maximum intensity projection shows multiple cells in the field of view spanning different axial planes. The lower right plot in each panel shows the z-profiles of cellular ROIs circled in the same colors on the image. The center of mass of each neuron ranges in depth from -5 to +4 µm.

Additionally, maximum intensity projections through *xz* and *yz* were generated with the TV term (Figure S1D). The TV term at both 10 and 30 iterations did not change the spatial signal confinement.

### 3.3 Light-field microscopy resolves calcium signals from neuronal dendrites in 3D

Light-field microscopy enables single-frame 3D imaging; therefore, we investigated its application to resolving calcium signals from neuronal processes in three spatial dimensions. We reconstructed 4D (x,y,z,t) light-field volumes from time series and extracted temporal signals from ROIs manually defined over the cell soma and two dendrites from the activation map.

Depth-time plots were extracted from ROIs taken from light-field time series reconstructed with synthetic refocusing (Figure 5B) and 3D deconvolution (3-iteration RL; Figure 5C). A depth map cannot be produced from widefield images focused on a single axial plane (Figure 5A).

**Fig 5.**
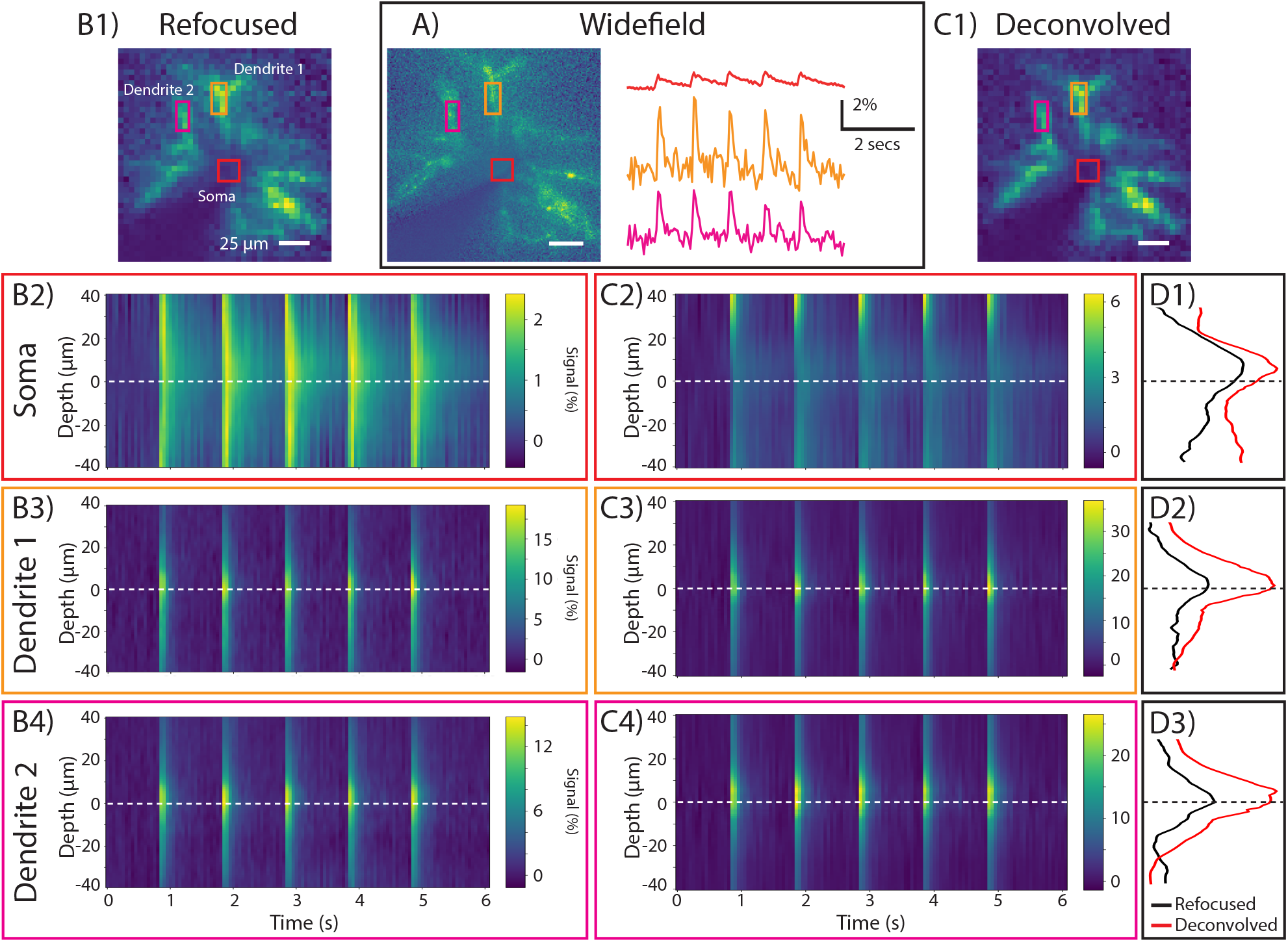
Calcium signals in dendrites can be observed across axially distinct planes from single-cell light-field volumes. A) shows the activation (or variance) map from a widefield image series with time courses extracted from a somatic ROI (red) and twol nearby dendrites (orange, pink). Depth-time plots are shown from the same ROIs reconstructed from a light-field time series with synthetic refocusing (B) and 3D deconvolution (C). D1-3 show the sum of the signal over time as a function of depth in the somatic and dendritic ROIs.

Somatic calcium transients can be seen across multiple planes in light-field time series reconstructed with synthetic refocusing (Figure 5B2) and 3D deconvolution (Figure 5C2). The signal as a function of depth has been summed over time (Figure 5D1). The somatic peak signal is greater in deconvolved volumetric light-field time series compared to those synthetically refocused, in agreement with the results from Section 3.

The increase in peak signal seen at the extremes of the axial range in the deconvolved light-field volumes is an artifact of the deconvolution algorithm and how the signal is calculated (eq. 5). The low baseline fluorescence and small dark signal is overpowered by the large out-of-focus dendritic fluorescent signal.

The peak signal seen in both of the dendrites is greater in deconvolved volumetric light-field time series (Figure 5C3,4) compared to those synthetically refocused (Figure 5B3,4), in agreement with the results from Section 3. From the depth plots it appears that the center of mass from both of the dendrite ROIs lie close to the native focal plane (∼5 µm) whereas the soma signal peaks at about 10 microns superficial to the native focal plane (Figure 5D2,3). This indicates that calcium transients can be resolved from neuronal subcompartments in axially distinct planes. Furthermore, the somatic signals spans a larger depths than the dendritic signals, corresponding the difference in their sizes.

The decay time, measured by the FWHM of somatic calcium transients at the native focal plane is the same between widefield (0.23 [0.20, 0.27]s, n=3 cells) and light-field time series reconstructed with synthetic refocusing (soma: 0.24 [0.21, 0.32]s, n=3 cells) and 3D deconvolution (soma: 0.22 [0.20, 0.40]s, n=3 cells). Moreover, there is no significant difference between the decay time of somatic and dendritic signals of synthetically refocused (dendrite: 0.139 [0.136, 0.141]s, n=3 cells) or deconvolved (dendrite: 0.132 [0.126, 0.167]s, n=3 cells) light-field time series.

## 4 Discussion

We resolved CaSiR-1 fluorescence transients in single cells and bulk-labeled live mouse brain slices. We found that calcium transient tSNR from bulk-labeled slices did not differ between widefield and light-field time series reconstructed with synthetic refocusing and three-iteration Richardson-Lucy 3D deconvolution. For single-labeled cells the tSNR was significantly larger for light-field time series reconstructed with three-iteration Richardson-Lucy 3D deconvolution compared to synthetic refocusing. Increasing the number of deconvolution iterations increased signal size and noise but reduced tSNR. Increased iteration number also increased axial confinement. Both light-field reconstruction algorithms, synthetic refocusing and Richardson-Lucy deconvolution, enabled 3D localization of calcium transients in single dye-loaded neurons and bulk-labeled slices. Extracting calcium transients from light fields, compared to widefield image time series, did not incur any penalty in terms of tSNR, while enabling volumetric imaging.

The reduction in SNR seen from deconvolved volumes arises from noise amplification due to lack of regularization.^29^ To reduce noise amplification, fewer iteration numbers provide a regularizing effect on the deconvolution.^13^ For higher iteration numbers, we attempted to overcome noise amplification by implementing TV-regularization in the RL deconvolution.^29^ However, this yielded no benefit in terms of signal, noise, or SNR in the extracted calcium time series.

Richardson-Lucy deconvolution at high iteration numbers decreases tSNR, and moreover increases computational cost compared to synthetic refocusing. In our implementation and hardware, reconstructing a volume (20 *µ*m) with synthetic refocusing took 40 seconds per frame while RL deconvolution took 20 seconds per iteration per frame (Processor i7 CPU @ 3.6 GHz, RAM 32 GB). A typical time series consisted of 200 frames (2048×2048 pixels, 20 Hz for 10 seconds). Reconstructing volumes (20 *µ*m) for the full time series took approximately 2 hours with synthetic refocusing, 3.5 hours 3-iteration RL deconvolution, and 22 hours with 20-iteration RL deconvolution. Methods to increase speed without the need to use high performance computing are desirable. Reconstruction speed has been improved by a number of groups through deep learning solutions.^19, 31^ However, the improved lateral and axial signal confinement achieved by iterative deconvolution methods may still motivate its use. We have shown that 3D deconvolution achieves higher spatial signal confinement than synthetic refocusing with axial confinement increasing at high iteration numbers. Therefore, to maximize spatial signal confinement a time-consuming iterative deconvolution technique could be beneficial.

Deconvolution algorithms leverage the fine sampling of individual projections through the volume, whereas refocusing cannot. Here we used a coarse deconvolution approach. Lateral oversampling can further improve the lateral signal confinement, providing lateral sampling rates greater than the native LFM resolution. However, oversampling increases computational cost and was unnecessary here as the LFM was designed for cellular resolution. We used the original light-field microscope design.^7^ Fourier light-field microscopy, where the microlens array is placed at the aperture stop of the microscope objective instead of the image plane, has also been shown to improve the lateral sampling rate even in the degenerate native focal plane.^32–34^

Both synthetic refocusing and 3D deconvolution reconstruction algorithms rely on ballistic photons, limiting their application in highly scattering mammalian brains. To minimize scattering, we used a red-emitting calcium dye, CaSiR-1 whose emission is less scattered than shorter wavelength emitting fluorophores. Furthermore, deep near-infrared indicators can be combined with blue-light sensitive opsins to achieve spectrally cross-talk free all-optical neurophysiology^35, 36^ or combined with shorter wavelength emitting fluorophores for imaging in multiple spectral channels.^37^ Nonetheless, scattering limited calcium signal extraction from reconstructed volumes to depths of approximately 50 microns, within the photon mean free path. Methods to improve signal extraction in scattering tissue have been demonstrated by computationally extracting fluorescence sources without reconstruction,^15, 20, 22, 38–40^ although reconstruction-less signal extraction cannot resolve the propagation of calcium signals throughout spatially extended structures such as dendrites. Combining the principles of confocal microscopy with LFM,^41^ selective-volume illumination,^19, 42, 43^ and/or spatially sparse labelling with genetically-encoded indicators can increase contrast to enable calcium signal extraction from reconstructed volumes at greater depths.

We detected dendritic calcium signals, evoked by back-propagating action potentials, in intracellularly dye loaded single cells. Limited dye diffusion precluded activity detection in distant processes. Applying LFM to neuronal tissues expressing genetically encoded calcium indicators (GECI) sparsely and strongly may enable tracing of functional signals through dendrites in three-dimensions, or synaptic mapping. Similar analyses have been performed for sparsely labeled genetically encoded voltage indicators (GEVIs) with a much lower baseline fluorescence, Δ*F/F*, and tSNR than that of the CaSIR-1 calcium dye.^44, 45^ Quicke *et al*. (2020) also demonstrated axial resolution of GEVI signals from dendrites at different depths. In combination with the present study, these results describe the LFM’s capacity to resolve function neuronal signals volumetrically at subcellular resolution in both low and high SNR regimes.

LFM captures 3D information with significantly reduced imaging time and bleaching compared to widefield. Generating similar 3D volumes in widefield would require physical refocusing of the objective in between trials. Our comparison of widefield trials to light-field trials reconstructed at the same axial plane revealed no penalty in terms of extracted calcium transient tSNR for light fields, which additionally enabled extraction of “in-focus” calcium transients from axially separated planes. Optically, implementing LFM is simple and low-cost, requiring only the introduction an off-the-shelf MLA at the native imaging plane of a standard widefield epifluorescence microscope. Cost-effective sCMOS cameras feature sensitivities and bandwidths well adapted to calcium LFM. Calcium imaging applications requiring high volume acquisition rates can readily benefit from LFM’s ability to trade spatial resolution for the ability to excite and image fluorescence simultaneously throughout a volume.

These results demonstrate the capabilities and limitations of two light-field reconstruction algorithms for high SNR calcium fluorescence imaging. The trade-offs described above highlight the importance of adapting the volume reconstruction strategy to the scientific goals and requirements of future neurophysiology experiments. For example, applications requiring online analysis to guide the experimental protocols would benefit from the speed and simplicity of synthetic refocusing or low iteration-number 3D RL deconvolution. We found that calcium signal extraction from volumes reconstructed with 3-iteration 3D RL deconvolution yielded high tSNR while bringing lateral signal confinement near to the maximum. However, higher iteration numbers, while decreasing tSNR, continued improving the axial confinement. These results demonstrate the importance characterizing and balancing tSNR, spatial signal confinement, and computational cost when selecting a volume reconstruction method for functional LFM applications.

## Acknowledgments

The authors would like to thank Yu Liu, Simon Schultz, and Ann Go for their technical assistance. The authors would also like to thank the Imperial College Research Computing Service. This project was funded by the Biotechnology and Biological Sciences Research Council (BB/R009007/1). National Institutes of Health (U01NS090501, U01NS099573, U01MH109091); Wellcome Trust Seed Award (201964/Z/16/Z); and a Royal Academy of Engineering Research Fellowship (RF1415/14/26).

## Author Contributions

CLH, PQ, and AJF conceived and designed the experiments. CLH and AJF designed the light-field optics. CLH performed experiments. CLH, PQ, and AJF designed the analysis. PQ, PS, HVJ, and PLD developed the deconvolution approach. CLH analyzed the data and wrote the paper. All authors contributed to manuscript revision and approved the final manuscript.

## Data, Materials, and Code Availability

The datasets and code generated for this study are available on request to the corresponding author.

**Carmel L. Howe** is a research associate at Imperial College London, UK. She received her MEng and Ph.D. degrees in Electrical and Electronic Engineering from the University of Nottingham in 2014 and 2018, respectively. She is currently developing a new high-speed, high-throughput, threedimensional imaging modality based on light-field microscopy to track network-level neuronal activity in the mammalian brain. Her research combines the fields of neurophysiology, optical engineering, signal and image processing.

**Peter Quicke** is postdoctoral research associate in the Department of Bioengineering at Imperial College London. He received his MSci. (with BSc.) degree in Physics in 2014, an MRes. in Neurotechnology in 2015 and his Ph.D. in 2019. His current research interests include computational microscopy and functional voltage imaging.

**Pingfan Song** is a research associate at Imperial College London. He obtained the Ph.D. degree at University College London (UCL), the master and bachelor degree both at Harbin Institute of Technology (HIT). His research interests lie in signal/image processing, machine learning and computational imaging with applications on a variety of image modalities.

**Pier Luigi Dragotti** received the Laurea degree (summa cum laude) in electronic engineering from University Federico II, Naples, Italy, in 1997, the master’s degree in communications systems from the Swiss Federal Institute of Technology of Lausanne (EPFL), Switzerland, in 1998, and the Ph.D. degree from EPFL, Switzerland, in April 2002. He has held several visiting positions. In particular, he was a visiting student at Stanford University, Stanford, CA, USA, in 1996, a Summer Researcher at the Mathematics of Communications Department, Bell Labs, Lucent Technologies, Murray Hill, NJ, USA, in 2000, and a Visiting Scientist at the Massachusetts Institute of Technology (MIT) in 2011. He is currently a Professor of signal processing with the Electrical and Electronic Engineering Department, Imperial College London. Before joining Imperial College in November 2002, he was a Senior Researcher at EPFL, working on distributed signal processing for sensor networks for the Swiss National Competence Center in Research on Mobile Information and Communication Systems. His research interests include sampling theory, wavelet theory and its applications, sparsity-driven signal processing with application in image super-resolution, neuroscience, and computational imaging. He was the Technical Co-Chair of the European Signal Processing Conference in 2012, an Associate Editor of the IEEE Transactions on Image Processing from 2006 to 2009, a member of the IEEE Image, Video and Multidimensional Signal Processing Technical Committee, and a member of the IEEE Signal Processing Theory and Methods Technical Committee. He was also a recipient of the ERC Investigator Award. He is currently the Editor-in-Chief of the IEEE Transactions on Signal Processing, and a member of the IEEE Computational Imaging Technical Committee and a Fellow of the IEEE.

**Amanda J. Foust** is a Royal Academy of Engineering Research Fellow and Lecturer in the Department of Bioengineering at Imperial College London. She studied Neuroscience with emphasis in computation and electrical engineering (BSc) at Washington State University, and Neuroscience (MPhil, PhD) at Yale University. The aim of her research programme is to engineer bridges between cutting-edge optical technologies and neuroscientists to acquire new, ground-breaking data on how brain circuits wire, process, and store information.

Biographies and photographs of the other authors are not available.

